# IRE1 RNase controls CD95-mediated cell death

**DOI:** 10.1101/2022.02.25.481813

**Authors:** D Pelizzari-Raymundo, R Pineau, A Papaioannou, XC Zhou, S Martin, T Avril, M Le Gallo, E Chevet, E Lafont

**Affiliations:** Inserm U1242, University of Rennes, Rennes, France; Centre de Lutte Contre le Cancer Eugène Marquis, Rennes, France

**Keywords:** CD95, Cell death, ER stress, Unfolded Protein Response, IRE1

## Abstract

Signalling by the Unfolded Protein Response (UPR) or by the Death Receptors (DR) represents cellular stress pathways frequently activated towards pro-tumoral outputs in cancer. Experimental evidence has highlighted functional links between the UPR and the DR TRAIL-R1/2. Herein, we demonstrate that the UPR sensor IRE1 controls the expression of CD95/Fas, another DR, and its cell death-inducing ability. Whereas CD95 is not a general determinant of ER stress-induced cell death, IRE1 RNase activity inhibition increased CD95 expression and exacerbated CD95L-induced cell death in glioblastoma (GB) and Triple-Negative Breast Cancer (TNBC) cell lines. In accordance, CD95 mRNA was identified as a target of Regulated IRE1-Dependent Decay of RNA (RIDD). Moreover, CD95 expression is elevated in TNBC and GB human tumours exhibiting low RIDD activity. Surprisingly, CD95 expression is also lower in XBP1s-low human tumour samples. We show that IRE1 RNase inhibition led to CD95 expression attenuation and reduced CD95-mediated hepatic toxicity in mice. In addition, overexpression of XBP1s increased CD95 expression and sensitized GB and TNBC cells to CD95L-induced cell death. Overall, these results demonstrate the tight IRE1-mediated control of CD95-dependent cell death signals in a dual manner through both RIDD and XBP1s, and they identify a novel, pharmacologically actionable link between IRE1 and CD95 signalling.

## Introduction

Throughout tumour development, cancer cells are subjected to both intrinsic and extrinsic stresses to which they must adapt to survive and proliferate. Adapted tumour cells survive this selection pressure through the activation of specific signalling pathways. Amongst those are the signalling engaged by the Death Receptors (DR) and the Unfolded Protein Response (UPR), an adaptive response to Endoplasmic Reticulum (ER) stress. The DR TRAIL-R1/2 and CD95 contribute to the immunosurveillance towards cancer or infected cells by their ability to induce cell death upon engagement by their ligand, TRAIL and CD95L, respectively ^1-3^. In some tumour cells however, engagement of CD95 or TRAIL-R1/2 fails to induce cell death and promotes pro-tumorigenic cellular outcomes. Similarly, constitutive UPR activation is often observed in cancer cells, resulting from adaptation to various stresses including oncogenic insults, aneuploidy or nutrient deprivation. This constitutive activation does not result in cell death, as would be the case in normal cells, but rather allows cancer cells to thrive through activation of pro-tumoral outcomes ^4, 5^. Thus, both UPR and DR signalling contribute to tumour progression and relapse in various cancer models including glioblastoma (GB) ^6-12^ and Triple Negative Breast Cancer (TNBC) ^13-16^. The UPR consists of three core signalling branches initiated by the activation of three sensors upon ER stress. These sensors are ATF6α (for Activating Transcription Factor 6 alpha), IRE1α (for Inositol-Requiring Enzyme 1 alpha, referred to as IRE1 hereafter) and PERK (for Protein kinase R-like ER Kinase). IRE1 RNase activity drives (i) the unconventional splicing of *XBP1* mRNA together with the tRNA ligase RtcB, ultimately leading to the expression of the transcription factor XBP1s, which in turn promotes the expression of multiple genes aimed at restoring ER homeostasis and (ii) the degradation of RNA through Regulated IRE1-Dependent Decay of RNA (RIDD). RIDD leads to both cytotoxic and non-cytotoxic cellular outcomes ^17^. In addition to their individual roles in controlling cell fate, both UPR and DR signalling pathways are functionally intertwined ^18, 19^. Indeed, TRAIL-R2, in some cases TRAIL-R1 and TRAIL, are upregulated downstream of the PERK/ATF4 and/or CHOP axis, in various ER stress conditions and accordingly TRAIL-R signalling can participate to cell death induced upon ER stress ^18-28^. On the contrary, RIDD limits TRAIL-R2-induced signalling by reducing the abundance of its mRNA ^22^. Moreover, TRAIL-R2 was recently shown to be directly activated intracellularly by misfolded proteins and therefore signals apoptosis from intracellular compartments^29^. Although the relationships between ER stress and TRAIL-R have been relatively well explored, whether and how IRE1 and CD95 signalling are also linked remains however, unclear. We have recently identified CD95 mRNA as being cleaved by IRE1 RNase in an *in vitro* RNA cleavage assay ^11^. Herein, we investigate the impact of IRE1 RNase activity in CD95 signalling, revealing a previously unrecognised dual functional link between these pathways.

## Results

### Basal and ER-stress induced IRE1 activation limits CD95 expression in GB and TNBC cells

We have recently identified CD95 mRNA as being cleaved by IRE1 in an *in vitro* RNA cleavage assay ^11^. To evaluate if the level of CD95 mRNA depends on IRE1 activity in a cellular context, we used the U87 GB cell line, which displays a constitutive activation of IRE1 ^11^. Expression of a dominant-negative (DN) form of IRE1, which represses the activation of endogenous IRE1, and thus its RNase activity ^30, 31^, led to a significant increase in the basal level of CD95 mRNA (**Fig 1A**). Accordingly, cell surface expression of CD95 was heightened in IRE1 DN-overexpressing U87 cells as compared to WT cells (**Fig 1B**). Similar results were obtained in two primary GB cell lines, RADH87 and RADH85 ^32^ in which expression of IRE1Q780*, a mutant of IRE1 devoid of both kinase and RNase domains that blunts IRE1 signalling, led to an increased expression of CD95 mRNA (**Fig 1A**) as well as of total and cell surface (**Fig S1A and Fig 1B**) CD95 protein. To determine if exogenous activation of IRE1 impacts on CD95 expression, we next analysed CD95 mRNA expression levels in U87 cells and in the TNBC cell line SUM159 treated or not with ER stress inducers. In this context, both tunicamycin (TM) and MG-132 prompted CD95 mRNA degradation, which was prevented by treatment with MKC-8866, a pharmacological inhibitor of IRE1 RNase activity ^33^ (**Fig S1B and C**). A consistent result was obtained regarding CD95 protein expression. Indeed, ER stress induced by TM or thapsigargin (TG) provoked a decrease in CD95 protein levels (**Fig 1C-F**) which was reverted by a treatment with MKC-8866. Overall, these results highlight that both constitutive and ER stress-induced IRE1 RNase activities limit CD95 expression in the tested cellular models.

**Figure 1.**
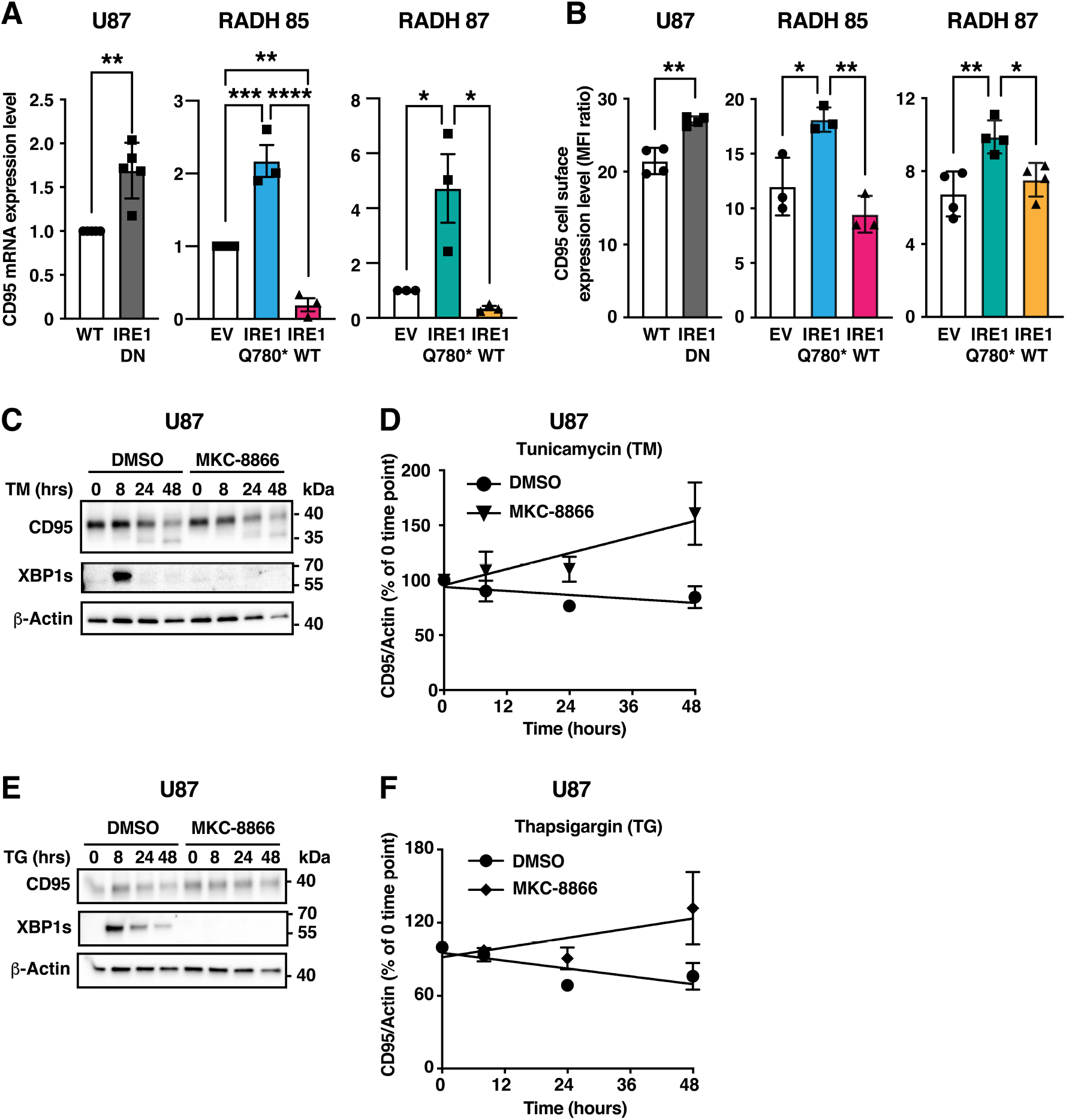
IRE1 regulates CD95 expression in TNBC and GB cells. **A**. mRNA was extracted from WT or IRE1 DN U87 cells and from empty vector (EV), IRE1WT- or IRE1Q780*-expressing RADH85 and RADH87 cells. CD95 mRNA was quantified by RT-qPCR and normalized to GAPDH. Mean ± SEM, n=3-5. **p < 0.01, *** p < 0.001, ****p < 0.0001 unpaired t-test with Welch’s correction (U87) or one-way ANOVA with Tukey multiple comparison correction for RADH85 and RADH87. **B**. CD95 cell surface level expression was evaluated by flow cytometry. Mean of MFI ratio ± SD, n=3. * p<0.05, **p < 0.01, unpaired t-test for U87; one-way ANOVA with Tukey multiple comparison correction for RADH85 and RADH87. **C, D** U87 cells pre-treated for 2 hours with MKC-8866 (30 μM) as indicated were further treated with 500 ng/mL tunicamycin for the indicated times. Lysates were analysed by western blot. **C**. One representative experiment out of three is shown. **D**. Quantification for three independent experiments is depicted. Mean ± SEM. **E, F**. U87 cells pre-treated for 2 hours with MKC-8866 (30 μM) as indicated were further treated with 50 nM thapsigargin for the indicated times. Lysates were analysed by western blot. **E**. One representative experiment out of three is shown. **F**. Quantification for three independent experiments is depicted. Mean ± SEM.

### IRE1 cleaves CD95 mRNA *in vitro*

To test whether CD95 mRNA is a target of RIDD, we first evaluated the ability of recombinant IRE1 to cleave CD95 mRNA *in vitro*. This revealed that CD95 mRNA is indeed directly targeted by IRE1 RNase activity (**Fig 2A**). We then searched the sequence of CD95 mRNA for the presence of potential RIDD cleavage sites. Such sites have been reported to both (i) reside in hairpin loop structures and (ii) display a consensus sequence CNG/CAGN^34-36^. However, no such site was identified in the CD95 mRNA sequence. We, therefore, expanded our search to additional potential sequences beyond this classical consensus. These supplementary sequences (CAACAA, CAGCUC, CUGCAU and CUGGCG) can be targeted by IRE1 *in vitro* when displayed on hairpin loops as recently identified in our laboratory ^37^. Based on this second analysis, we identified two potential cleavage sites, one located within the CD95 ORF and the other in the 3’UTR of CD95 mRNA (**Fig 2B**). Noteworthy, PCR amplification of 136 bp and 121 bp regions encompassing the CAACAA and CUGCAU sites, respectively, was dampened when performed using RNA incubated with recombinant IRE1, arguing that these two sites might indeed be targeted by IRE1 *in vitro* (**Fig 2C, D)**. Taken together, these results indicate that CD95 mRNA is a *bona fide* substrate of IRE1 RNase *in vitro*.

**Figure 2.**
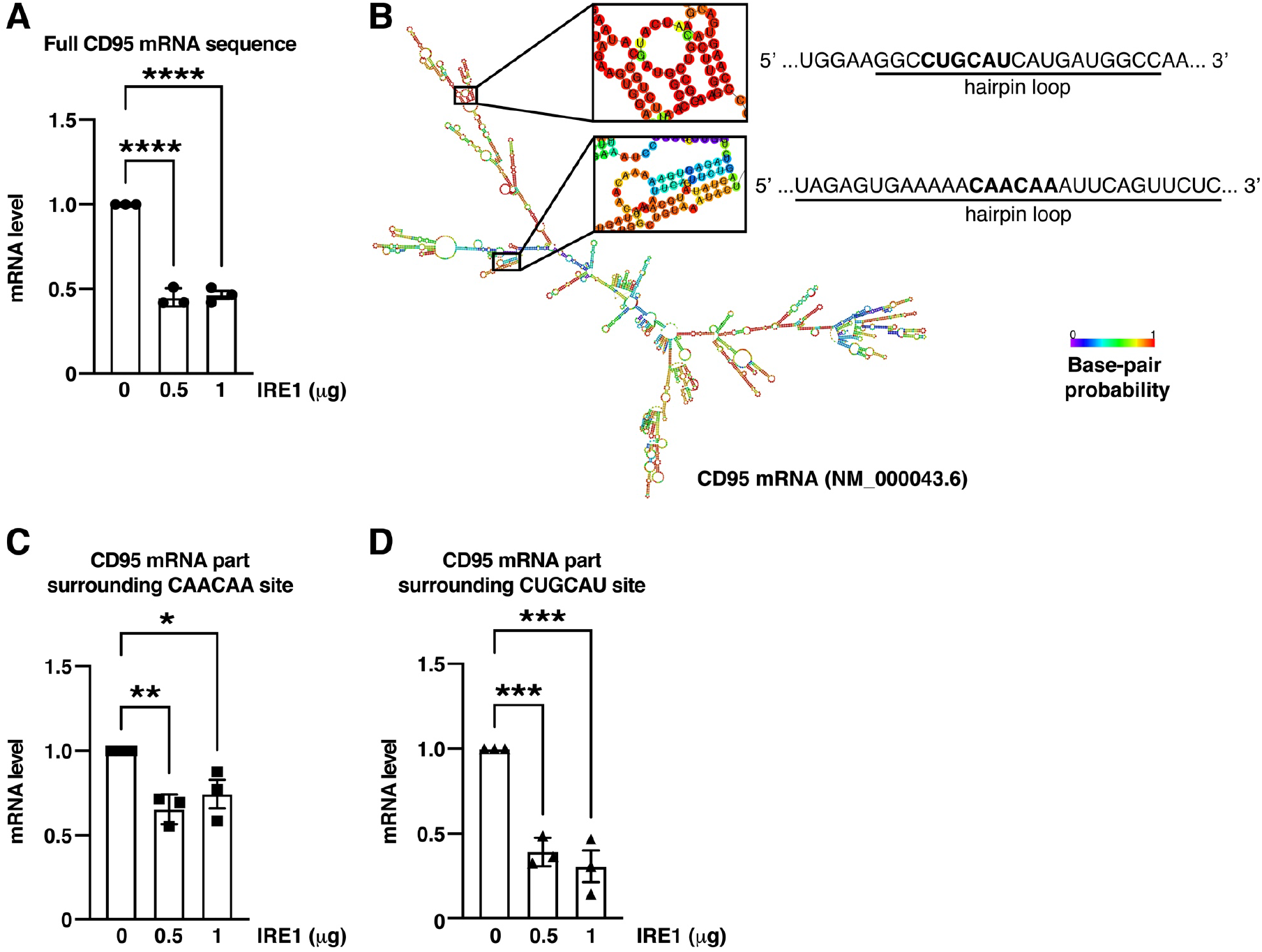
IRE1 cleaves CD95 mRNA *in vitro*. **A**. RNA (2 μg) extracted from U87 cells was incubated with the indicated amounts of recombinant IRE1 for 1 hour. CD95 mRNA was then quantified by RT-qPCR and normalized to GAPDH. Mean ± SEM, n=3. One-way ANOVA with Dunnett multiple comparison correction, ****p < 0.0001. **B**. Predicted folded structure of CD95 mRNA. The two predicted cleavage sites within hairpin loops are highlighted. **C, D**. RNA (2 μg) extracted from U87 cells was incubated with the indicated amounts of recombinant IRE1 for 1 hour. 136-bp (C) and 121-bp (D) parts of CD95 mRNA surrounding the indicated potential cleavage sites were then quantified by RT-qPCR and normalized to GAPDH. Mean ± SEM, n=3. One-way ANOVA with Dunnett multiple comparison correction, *p < 0.05, **p < 0.01, *** p < 0.001.

### CD95 is not a general determinant of ER stress-induced cell death

Since the expression of another DR, TRAIL-R2, is also regulated by IRE1 ^22^ and that TRAIL-R2 signalling impacts on ER stress-induced cell death, we next hypothesized that CD95 may also influence the sensitivity of cells to ER stress. To test this hypothesis, we used U87 cells in which CD95 was knocked-down using RNA interference as well as MDA-MB-231 TNBC cells WT or KO for CD95 (**Fig S2A, B**). Whereas CD95 depletion led to a reduced sensitivity to TM-induced loss of viability in both cell lines, it did not impact any of these cells’ ability to die in response to TG, MG-132 and Brefeldin A (**Fig 3A, B and Fig S2A, B**). These results, therefore, imply that while CD95 might contribute to cell death in response to some specific ER homeostasis insults or to aberrant protein N-linked glycosylation, this DR is not a universal determinant of cells’ sensitivity to ER stress-induced cytotoxicity.

**Figure 3.**
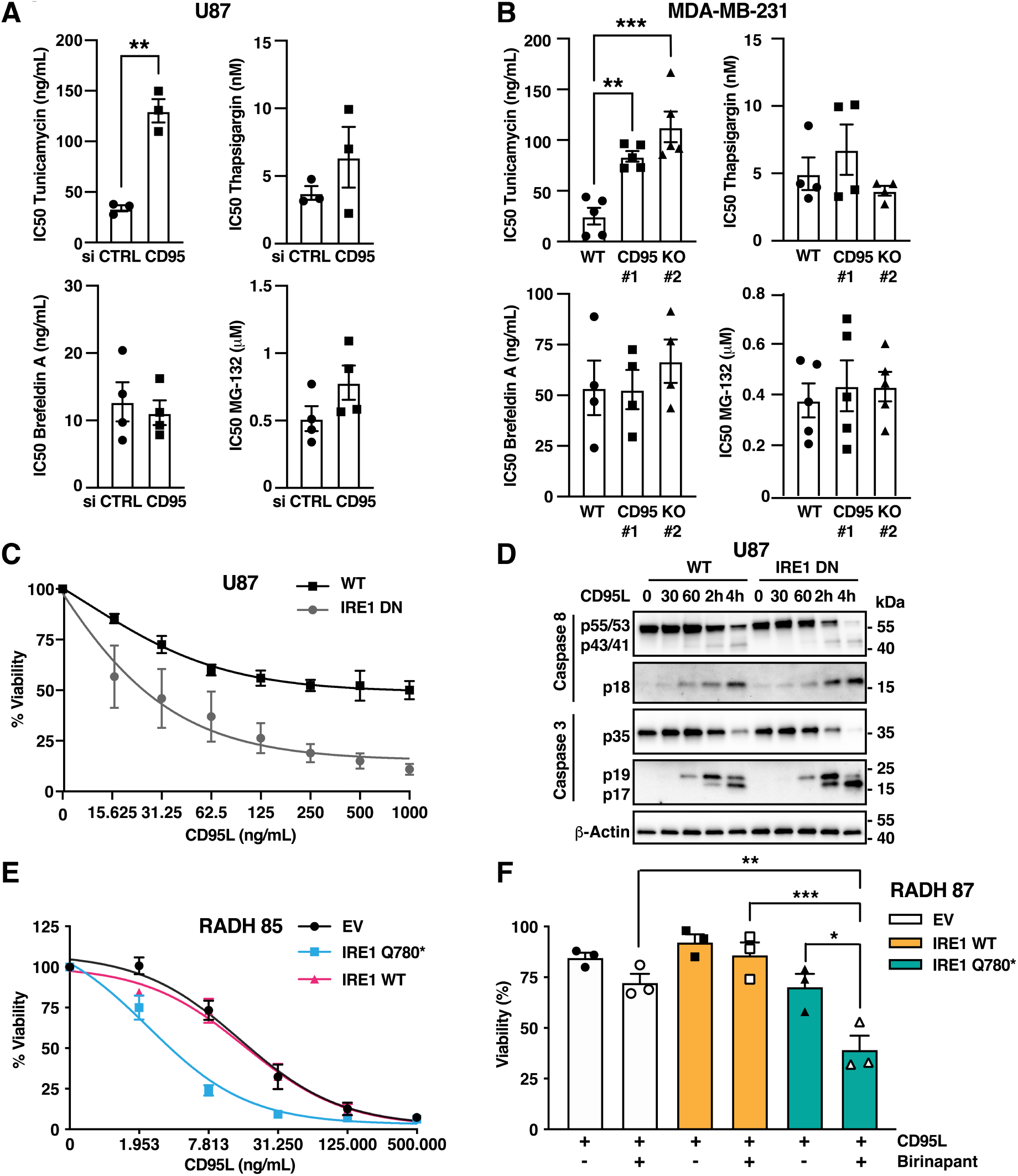
CD95 is not a universal determinant of ER-stress induced cell death whilst IRE1 RNase activity limits CD95L-induced cell death. **A**. U87 were transfected with siRNA control or targeting CD95. 48 hours later, cells were treated for 48 hours with the indicated ER stress inducers. Viability was determined using an MTT assay. The relative IC50 calculated for each independent experiment is represented (see also Fig S2A). **p < 0.01, unpaired t-test **B**. MDA-MB-231 WT or CD95 KO clones were treated for 48 hours with the indicated ER stress inducers. Viability was determined using an MTT assay and relative IC50 calculated for each independent experiment (see also Fig S2B). **p < 0.01, *** p<0.001, one-way ANOVA with Tukey multiple comparison correction **C**. U87 WT or expressing IRE1DN were treated with the indicated concentrations of CD95L for 24 hours. Viability was determined using MTT assay and normalized to untreated cell values. Mean ± SEM of 3 independent experiments. **D**. U87 WT or expressing IRE1DN were treated with 250 ng/mL CD95L for the indicated times. Lysates were analysed by western blot. One experiment representative of three independent ones is shown. **E**. RADH85 control (EV), stably expressing IRE1Q780* or IRE1WT were treated with the indicated concentrations of CD95L for 24 hours. Viability was determined using MTT assay and normalised to untreated cells values. Mean ± SEM of 3 independent experiments. **F**. RADH87 control (EV), stably expressing IRE1Q780* or IRE1WT were pre-treated with 50 nM Birinapant and further treated with 1 μg/mL CD95L for 24 hours. Viability was determined using MTT assay and normalised to untreated cells values. Mean ± SEM of 3 independent experiments. **p<0.05, **p < 0.01, ***p < 0.001, one-way ANOVA with Tukey multiple comparison correction.

### IRE1 RNase activity limits CD95L-induced caspase-activation and cell death

Since our cell-based results indicate that RIDD limits CD95 expression, we hypothesized that IRE1-mediated depletion of CD95 would repress CD95L-induced cell death. Indeed, U87 cells expressing IRE1 DN were markedly sensitized to CD95L-induced loss of viability (**Fig 3C**). In line with this observation, activation of the initiator caspase-8 and the effector caspase-3 happened earlier and was more pronounced in IRE1 DN-overexpressing U87 cells as compared to WT cells (**Fig 3D**), further pointing out that IRE1 controls an early event in CD95 cell death signalling. Noteworthy, RADH85 cells expressing IRE1Q780* were also sensitized to CD95L-induced cell death (**Fig 3E**). Yet, despite increased expression of CD95 (**Fig 1B**), RADH87 expressing IRE1Q780* were not sensitized to CD95L-induced cell death (**Fig 3F**), leading us to postulate that an additional cell death-inhibitory checkpoint, downstream of CD95, might be active in these cells. To test this hypothesis, we targeted the anti-apoptotic proteins cIAP1/2 and XIAP which are known negative regulators of CD95-mediated cell death ^38, 39^. To this end, we utilized the SMAC mimetic Birinapant which potently inhibits cIAP1 and, albeit to a lesser extent, cIAP2 and XIAP ^40, 41^. RADH87 expressing IRE1Q780^*^ were notably sensitized to CD95L-induced loss of viability when co-incubated with Birinapant as compared to control or IRE1 WT-overexpressing cells (**Fig 3F**). Taken together, these data argue for an early role of IRE1 in limiting CD95L-induced cell death signalling.

### Impact of IRE1 RNase inhibition in CD95-mediated cell death in an acute liver injury mouse model

To determine whether IRE1 RNase activity controls CD95 expression and signalling *in vivo*, we first evaluated the impact of intra-peritoneal injection of MKC-8866 on CD95 expression in mouse livers (**Fig 4A**). In contrast with the results previously obtained in cancer cell lines (**Fig 1-2**), this analysis indicated that the pharmacological inhibition of IRE1 RNase led to decreased CD95 protein expression *in vivo* (**Fig 4 B**). To investigate whether this unexpected reduction of CD95 expression levels coincided with a decreased sensitivity to CD95-mediated cell death, we next used the anti-CD95 agonist antibody Jo2 at a sub-lethal dose, known to induce a mild hepatitis within 6 hours of treatment ^42^. Mice pre-treated with MKC-8866 or vehicle were therefore injected with Jo2 or the corresponding isotype control (**Fig S3A**). First, immunohistochemical (IHC) analysis of the liver sections confirmed that inhibition of IRE1 led to a reduction in CD95 expression (**Fig S3B**). Furthermore, both HES and IHC analysis of the same liver sections using an anti-cleaved caspase-3 indicated a clear reduction of Jo2-induced liver damage in the MKC-8866-treated group as compared to vehicle-treated animals (**Fig 4C and Fig S3C**), a result which was further confirmed by Western blot (**Fig S3D**). Taken together, these results indicate that IRE1 RNase activity promotes CD95 hepatic expression and CD95-mediated hepatotoxicity *in vivo*.

**Figure 4.**
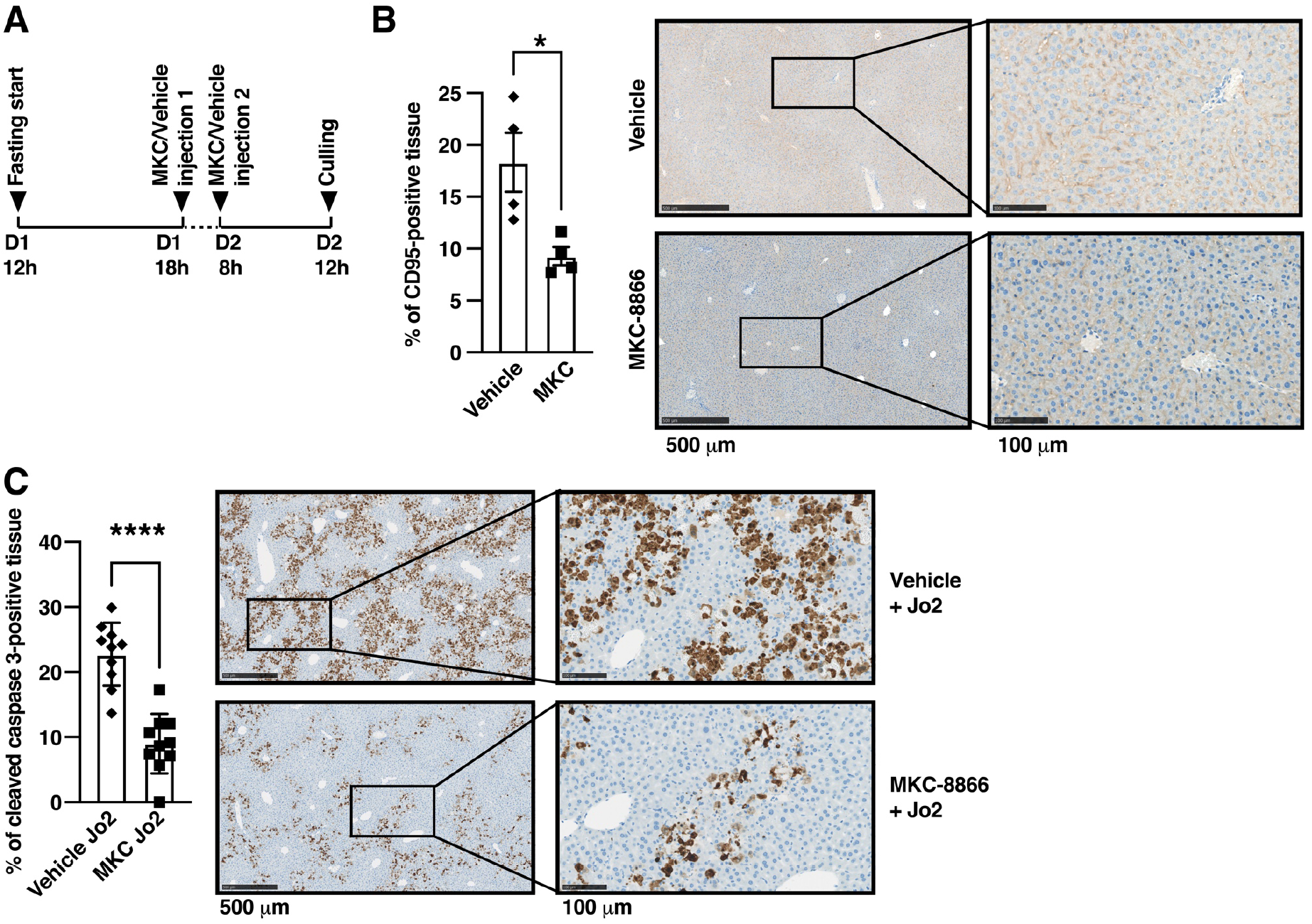
IRE1 RNase inhibition limits hepatic CD95 expression and CD95-mediated cell death in mice. **A**. Timeline of *in vivo* experiment 1. Eight mice were divided in two groups of 4 and repeatedly injected as indicated with either vehicle (group 1) or MKC-8866 (group 2). **B**. IHC staining for CD95 on liver tissue from the two groups of mice described in A. Left: quantification of CD95 staining. Mean ± SEM of n=4 mice per group; *p<0.05 with Mann-Whitney test for comparison of the two groups. Right: representative IHC image for each group. **D**. IHC staining for cleaved caspase-3 on liver tissue from the two indicated groups of mice described in Fig S3A. Left: quantification of cleaved caspase-3 staining. Mean ± SEM of n= 10 mice per group; ****p<0.0001 Mann-Whitney test for comparison of the two indicated groups. Right: representative IHC image for each group.

### Dual regulation of CD95 expression and signalling by IRE1 RNase activity

To further investigate the apparent discrepancy in the results obtained in cell lines and *in vivo*, we explored how the different branches of IRE1 signalling impact on CD95 expression. Indeed, IRE1 RNase activity catalyzes both RIDD and the unconventional splicing of XBP1 mRNA, the latter resulting in the expression of the XBP1s transcription factor. Since we observed that MKC-8866 dampened CD95 expression and CD95-mediated cell death *in vivo*, we thus hypothesized that XBP1s, contrary to RIDD, may enhance CD95-mediated cell death. This hypothesis was validated as we observed that overexpression of XBP1s in both U87 and SUM159 increased CD95L-induced cell death **(Fig 5A, B)**. Next, we evaluated the impact of XBP1s overexpression on the protein level of pro-apoptotic components of the CD95 pathway in SUM159 or U87 cells. Amongst these factors, solely CD95 expression was induced by high XBP1s overexpression in both cell lines (**Fig 5C**). This therefore suggests that CD95 could be a genuine XBP1s-target gene. It remains to be evaluated whether this transcription factor sensitizes cells solely through up-regulation of this DR, or of yet-to-be identified pro-apoptotic factors of this pathway, a combination thereof or additional more indirect mechanisms. Nevertheless, together with the *in vivo* data (**Fig 4**), our results point towards a role for XBP1s in promoting CD95-mediated cell death. Collectively our results indicate a dual regulation of CD95 expression and signalling by IRE1 RNase activity. Next, we explored whether the activation of RIDD and XBP1s might correlate with CD95 mRNA expression levels in human tumours. Thus, we analysed the expression of this mRNA in tumours classified according to their RIDD and XBP1s gene expression signature, as defined previously^11^. This analysis revealed that XBP1s-high tumours present a significantly heightened expression of CD95, as compared to XBP1s-low tumours (**Fig 5D**). Conversely, RIDD-high tumours present a significantly diminished CD95 mRNA level when compared to RIDD-low tumours (**Fig 5D**), hence indicating a dual regulation of CD95 mRNA expression by IRE1 signalling. A broader analysis of the expression of other DR (TNFR1, TRAIL-R1, TRAIL-R2) as well as the initiators of apoptosis FADD and caspase-8 revealed that most of these factors were also significantly differently expressed dependently on the XBP1s and RIDD signature (**Fig S4 and S5**). Intriguingly, the expression of these components seemed to be overall preferentially correlated with the activation of XBP1s in TNBC tumours, whilst being overall negatively correlated with RIDD in GB tumours (**Fig S4 and S5**).

**Figure 5.**
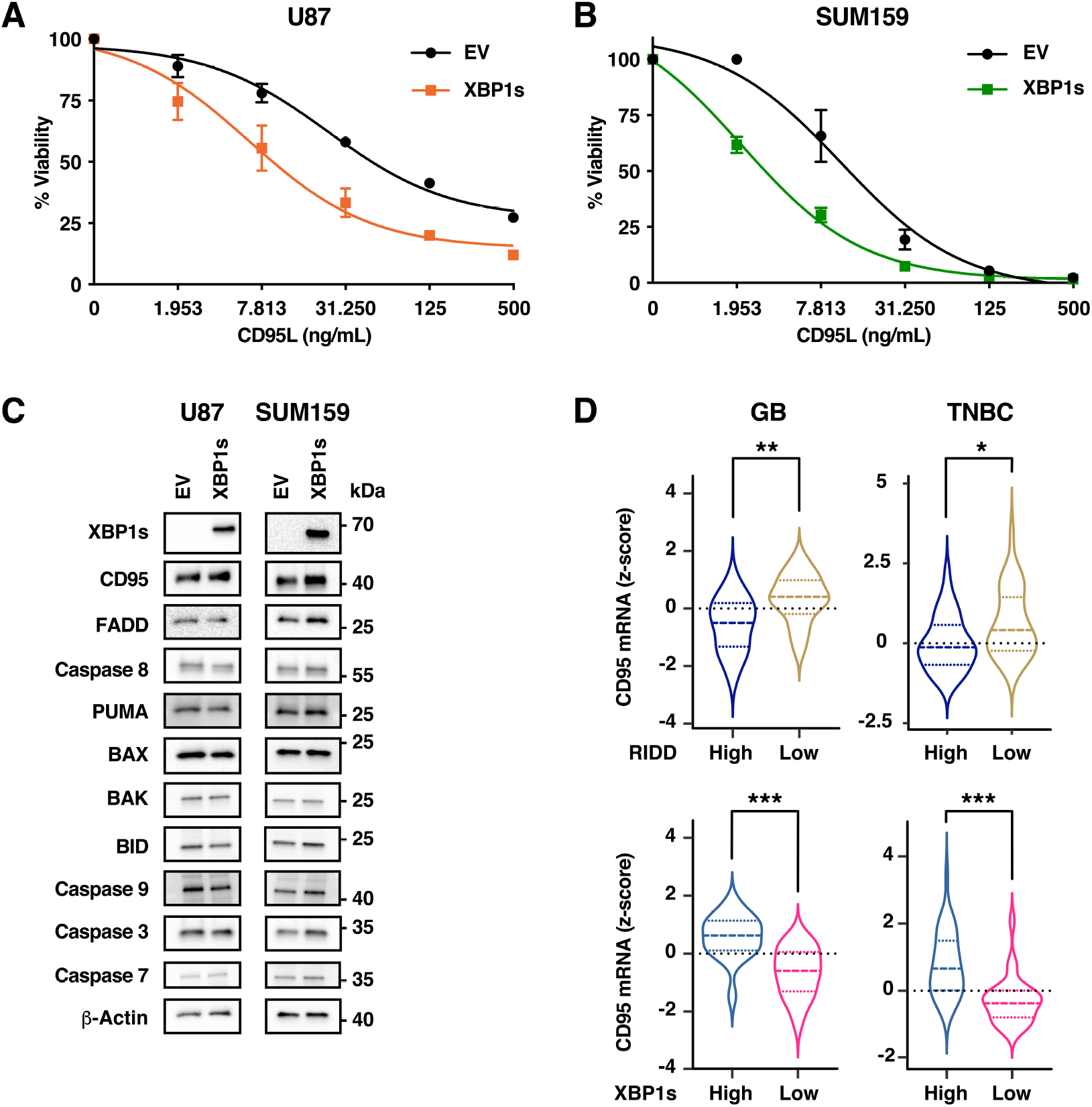
XBP1s promotes CD95L-induced death and the relative activation of each IRE1 branch correlates with CD95 expression in tumors. **A**. U87 cells were transfected with a plasmid coding for FLAG-XBP1s (XBP1s) or an empty vector (EV). 48 hours later, cells were treated with the indicated concentrations of CD95L for 48 hours. Viability was assessed using MTT assay and normalized to untreated cell values. Mean ± SEM of 3 independent experiments. **B**. SUM159 cells were transfected with a plasmid coding for FLAG-XBP1s (XBP1s) or an empty vector (EV). 48 hours later, cells were treated with the indicated concentrations of CD95L for 48 hours. Viability was assessed using MTT assay and normalized to untreated cell values. Mean ± SEM of 3 independent experiments. **C**. U87 or SUM159 cells were transfected with a plasmid coding for FLAG-XBP1s (XBP1s) or an empty vector (EV). 48 hours later, cells were lysed and lysates were analysed using western blot. **D**. CD95 expression z-scores of 45 GB and 67 TNBC tumors were plotted according to the RIDD activity score (top) and according to the XBP1s activity score (bottom). The distribution of z-score is represented as violin plots. Statistical difference of expression between groups was calculated using Mann-Whitney tests and the p-value is indicated (** p<0.01, **** p ≤0.0001).

## Discussion

Both UPR and DR signalling contribute to the elimination of cancer cells whereas diversion of these pathways towards pro-tumorigenic outcomes can promote tumour progression. So far, extensive experimental evidence demonstrated that TRAILR-1/2-signalling and the UPR are functionally linked, via PERK and IRE1 through its RIDD activity, to control cell fate^18-28, 43^. TRAIL-R1/2-emanating signals not only contribute to ER stress-induced cell death but also to cytokine production^44^. Furthermore, it has recently been shown that TRAIL-R2 can directly bind to and get activated by misfolded proteins^29^ to promote cell death from intracellular compartments^45^. Of note, the TRAIL-R1 gene has also been identified as a target of XBP1s in a chromatin immunoprecipitation experiment, and it is thus tempting to speculate that the expression of this DR might be directly inducible by this transcription factor at least in some cell types^13^. With regards to CD95, it was previously reported that its expression can be induced by ER stress in macrophages, a phenomenon which was attributed to a calcium/calmodulin-dependent protein kinase IIγ (CaMKIIγ)-dependent pathway^46^. In addition, in INS-1E rat insulinoma cells, it was suggested that cyclopiazonic acid could enhance IL-1-induced CD95 expression in an XBP1s-dependent manner^47^. Therefore, the existence of a potential connection between ER homeostasis control and CD95 in cancer cells remained largely uncharted. Herein, we observe that IRE1 RNase cleaves CD95 mRNA *in vitro* and IRE1 activity represses CD95 expression (at both mRNA and protein levels) as well as CD95L-induced cell death in TNBC and GB cell lines. Accordingly, CD95 mRNA expression is reduced in both GB and TNBC human tumours displaying an elevated RIDD activity. Unexpectedly, we observed that expression of CD95 mRNA is increased in XBP1s-high tumours. This observation coincided with the fact that overexpression of XBP1s increases CD95 expression, sensitizes cells to CD95L-induced cell death and that pharmacological inhibition of IRE1 RNase represses both CD95 hepatic expression and the hepatotoxicity induced by a CD95-targeting agonistic antibody. Therefore, our study highlights a previously unrecognised and dual link between IRE1 activity and the expression and signalling of CD95 (**Fig 5E**).

The pathophysiological importance of such connection remains to be fully explored, but given the increasing variety of physiological and pathological conditions influenced by CD95 and/or IRE1^3, 4^, we deem it likely to be broadly relevant. One such context could be tumour progression. Indeed, the ability of IRE1 to dually regulate CD95-mediated cell death suggests that the preferential branch activated by this RNase in cancer cells could be one of the determinants of their response to endogenous immunosurveillance or even T-cell immunotherapy^48, 49^. Furthermore, this study and previous work^22^ suggests that the expression of additional DR along that of initiator caspase-8 and FADD by tumour cells could be controlled by IRE1. Thus, one could speculate that such a regulation could contribute to the anti-tumoral effect of IRE1 overexpression recently reported in immunocompetent mice^50^. CD95 can also mediate a variety of non-cytotoxic pro-tumoural cellular outcomes, including migration, production of pro-inflammatory cytokines, proliferation and regulation of cell differentiation state, which are also modulated by IRE1 signaling^2, 3, 51^. Therefore, it is likely that IRE1-mediated CD95 increased expression could contribute to tumour progression in cancer cells which display primary or acquired resistance to CD95L-induced cell death. Hence, to promote CD95L-induced cell death over non-apoptotic cellular outcomes, it will also be required to define tumour contexts in which additional cell death checkpoints (*e*.*g*. mediated by IAPs) should be alleviated in concert with IRE1 targeting. Additional pathological conditions have been shown to be potentially regulated by both CD95 and IRE1. For example, IRE1 inhibition prevents fibrosis in an idiopathic pulmonary fibrosis (IPF) murine model^52, 53^. In this context, loss of CD95 promotes persistent fibrosis^54^, thus pointing towards this non-cancer model to also evaluate the IRE1/CD95 relationship.

Our data indicate that CD95-dependent cell death signals are oppositely regulated by RIDD and XBP1s. Such a phenomenon could therefore represent a mechanism to fine-tune life and death decisions. In addition, such dual mechanism might not be a rare occurrence, as suggested by our observation that the expression of additional DR signalling components correlates with the relative activation of XBP1s and/or RIDD in GB and TNBC. Furthermore, ongoing work from the laboratory suggests that more than 10% of XBP1s target genes may also be potential RIDD substrates^6^. Whilst in the context of CD95 and other DR signalling this dual regulation may be an additional way to ensure timely cell death induction, it will be interesting to explore the functional consequences of modulating the other yet-to-be-validated dual targets of IRE1 signalling. Such a knowledge will also help to define the specific tumour contexts in which pharmacological inhibition or activation of IRE1 may be beneficial or detrimental.

## Material and methods

### Cell lines, cell culture and reagents

U87, RADH85 and RADH87 (WT, DN, EV, IRE1WT or IRE1Q780*) and MDA-MB-231 WT or CD95 KO1 and 2 (initially named CD95 KO 5 and KO 9 respectively) were generated in our laboratory as described previously^11, 30-32, 55^. All cell lines were cultured in Dulbecco’s modified Eagle’s medium (DMEM) supplemented with 10% decomplemented FBS and 2 mM L-glutamine at 37°C in a 5% CO_2_ incubator. Modified RADH85 and 87 were cultured with 0.8 μg/mL or 1 μg/mL puromycin, respectively. All cells were regularly tested for mycoplasma absence. CD95L was produced and quantified in-house as described previously^56^. Thapsigargin (SML1845), Actinomycin D (A9415) and Tunicamycin (T7765) were from Sigma-Aldrich. Brefeldin A (S7046), MKC-8866 (S8875), Birinapant (S7015), MG-132 (S2619) were from Selleckchem.

### RT-qPCR

Total RNA was extracted from cells using Trizol (Thermo-Fisher Scientific, Thermo-Fisher Scientific, 15596026) according to the manufacturer’s instructions. cDNA was synthesized from the total RNA using the Maxima Reverse Transcriptase enzyme, random hexamer primers, dNTP mix and the Ribolock RNase inhibitor (Thermo-Fisher Scientific). PCR was performed on the template cDNA using Phusion High-Fidelity DNA Polymerase and dNTP mix (Thermo-Fisher Scientific). Quantitative PCR was alternatively performed for the cDNA using the SYBR® Premix Ex Taq™ (Tli RNase H Plus) (TAKARA-Clontech) using a QuantStudio5 system (Applied Biosystems). The primer sequences used for these experiments are shown in Table 1.

**Table 1.**
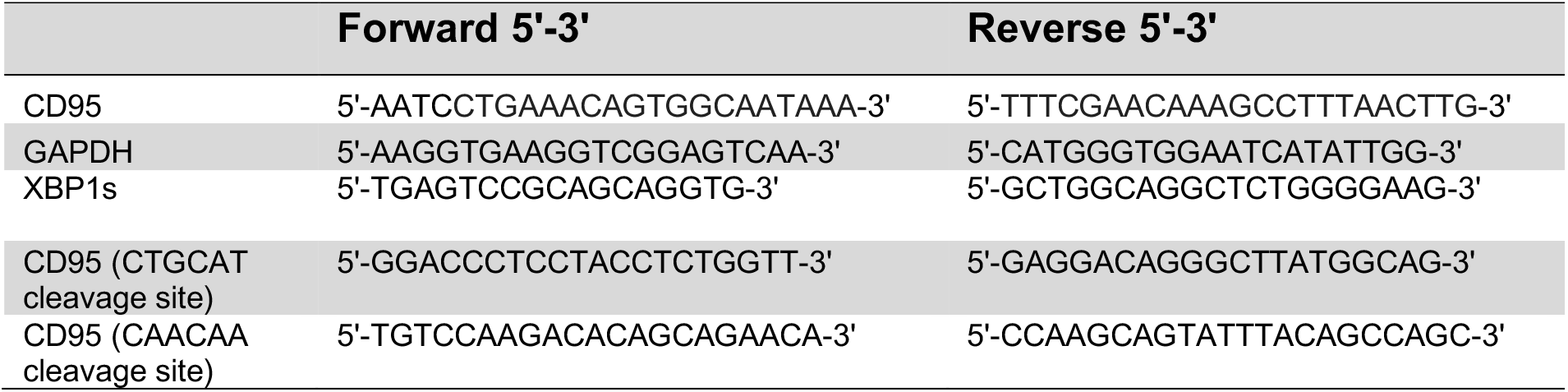
RT-qPCR primers to monitor the expression of Homo sapiens CD95, XBP1s and GAPDH mRNA

### In vitro RNA cleavage assay

The predicted CD95 mRNA structure was obtained using the RNAfold Web server ^57^. Different concentrations of recombinant IRE1(Sino Biological) were incubated with 2 μg of RNA extracts from U87 cell line, 2mM DTT (Merck, D3801), 20 mM ATP (Merck, A2383) in 20mM Tris, 500mM NaCl, 10% gly, pH7.4 for 1 hour at 37°C. After the incubation, the resulting samples were used to perform qRT-PCR as described above.

### Cell and tissue lysis and western blot

For cell lines: cells seeded in 6 well plates were treated as indicated in the figure legends and lysed in 200-250 μL ice-cold RIPA buffer (Tris-HCl 50 mM, ph7.4, NaCl 150 mM, EDTA 2 mM, Sodium Deoxycholate 0.5%, Triton 1% ; SDS 0.1%) per well, including proteases (Protease inhibitor cocktail, Sigma-Aldrich, P8340) and phosphatase inhibitors (Phosphatase inhibitor cocktail 2, Sigma-Aldrich, P5726). After 10 minutes incubation on ice, lysates were briefly sonicated and cleared by centrifugation (15500g, 30 min, 4°C). Lysates were collected and protein quantified using Pierce™ BCA protein assay (Thermo scientific). Laemmli sample buffer (BIO-RAD, 1610747) with 2-mercaptoethanol (BIO-RAD, 1610710) was added to the samples which were heated (95°C, 5 min). For liver tissue: tissue pieces (300-450 mg) were flash-frozen using liquid nitrogen upon collection. Tissues were homogenized with a Precellys Lysing kit (Bertin Ref. P000911) according to the manufacturer’s instructions. For western blot analyses of cell and tissue lysates, 10-50 μg protein was loaded on 10% acrylamide home-made or gradient (4-15% acrylamide) commercial (BIO-RAD, 4561086) Tris-Glycine-SDS gels along a protein ladder (PageRuler prestained protein ladder, Thermo scientific, 26617). Proteins were then transferred onto nitrocellulose membranes using the turbo transfer system (BIO-RAD). Membranes were saturated with TBS with 0.5% Tween-20 and 5% Milk for 1 hour prior to overnight 4°C incubation with the indicated primary antibody diluted 1/1000° in TBS-Tween-5%BSA 0.025% Sodium Azide. HRP-coupled secondary antibodies, used at 1/5000° in TBS-Tween-5% milk, were from Southern Biotech. Signal Fire™ ECL reagent (Cell signaling) or ECL Revelblot intense (OZYme, OZYB002-1000) was used and chemiluminescence signal was detected using G:Box Chemi XX6 imager from Syngene. For quantification, the Image J software was used^58^. The following primary antibodies were used for western blot: CD95 (Cell signaling, 4233), Actin (Sigma-Aldrich, A5316), IRE1 (Cell signaling, 3294 and Santa Cruz, 390960), XBP1s (Biolegend, 647502), Caspase-8 (Adipogen, AG-20B-0057-C100, Caspase-3 (Cell signaling, 14220), Cleaved caspase-3 (Cell signaling, 9661), FADD (Cell signaling, 2732), PUMA (Cell signaling, 12450), BAX (Cell signaling, 5023), BAK (Cell signaling, 12105), BID (Cell signaling, 2002), Caspase-7 (Cell signaling, 12827), Caspase-9 (Cell signaling, 9502).

### Flow cytometry

To evaluate CD95 expression at the cell surface, cells (U87, RADH85 or RADH87) were seeded in 6 well plates (250000 cells/well). 24 hours later, cells were harvested and stained using Zombie-violet™ (Biolegend, 423113) diluted in PBS according to the manufacturer protocol. Then, samples were saturated with PBS containing 1% BSA and 1% FCS and human FcR-blocking reagent (Miltenyi Biotec, 130-059-901) for 15 minutes at 4°C prior to labelling using an anti-human CD95-APC antibody (clone DX2, Miltenyi Biotec, 130-117-701) or corresponding IgG1-APC isotype (clone IS5-21F5, Miltenyi Biotec, 130-113-196) for 30 minutes at 4°C. Following two washes with the saturation buffer, cells were resuspended in PBS and analysed using a Novocyte cytometer (Acea Biosciences). CD95 cell surface expression was analysed after gating on viable singlets. Results are represented as a ratio of Median Intensity Fluorescence values for anti-CD95-stained over isotype-stained samples.

### Transfection

The cells indicated in the legend were seeded in 6 well plates at 250000 cells per well. For knockdown experiments, cells were transfected 24 hours after seeding using LipoRNAimax (Invitrogen, 13778075) according to the manufacturer protocol with 80 nM siRNA (On-target plus Non-targeting or Human FAS pool, Horizon Discovery, D-001810-10-05 or L-003776-00-0005, respectively). 24 hours after transfection, cells were reseeded in 48 well plate and 6 well plate for viability and western blot analysis respectively. 24 hours after reseeding, cells were either lysed (for western blot) or treated as described for viability assay. For over-expression experiments, cells were transfected 24 hours after seeding using 2.5 μg DNA per well and lipofectamine 3000 (Invitrogen, L3000008) according to the manufacturer protocol. pcDNA3.1+-XBP1s-FLAG was obtained from Genescript (OHu25513D Accession No: NM_001079539.1, Vector: pcDNA3.1+/C-(K)DYK Species: Human). 24 hours after transfection, cells were reseeded in 48 well plate. 24 hours after reseeding, cells were treated as described for viability assay.

### Viability assay

Cells seeded in 48 well plates (20000 cells per well) were treated as indicated in the figure legends. 500 μg/mL MTT (Invitrogen, M6494) was added to the medium for the last two hours of treatment. Medium was then removed and the formazan crystals were solubilized using DMSO. Reading of the absorbance was performed at 560 nm using a Tecan Infinite F200 Pro reader. Results are expressed as % of viability, with 100% being untreated cells for each cell line or experimental condition.

### In vivo experiments

All in vivo experiments described in this study have been approved by the “Comité d’éthique de l’Université de Rennes 1” and the French ministry of education, research and innovation under the licence APAFIS #32241-2021061714545440. These experiments were performed at the ARCHE-BIOSIT UMS 34380 (Rennes). 8-weeks old C57BL/6rJ male mice were obtained from Janvier. Mice were fasted on day 1 at 12 pm for both experiments. Volumes injected intra-peritoneally did not exceed 250 μL for both experiments. For experiment 1, intra-peritoneal injections of MKC-8866 (or equivalent volume of DMSO) diluted in PBS to 1 mg/mL were performed at 6 pm on day 1 and 8 am on day 2 (10 mg/kg per injection) and mice were culled at 12 pm on day 2. For IHC analyses, a liver piece (around 200 mg) was fixed in paraformaldehyde 4% solution. The rest of the liver was rapidly cut in small pieces prior to flash-freezing using liquid nitrogen for western blot analysis. For experiment 2, intra-peritoneal injections of MKC-8866 (or equivalent volume of DMSO) diluted in PBS to 1 mg/mL were performed at 12 pm on day 1, 6 pm on day 1 and 8 am on day 2 (10 mg/kg per injection). At 12 pm on day 2, an intra-peritoneal injection of 0,15 mg/kg of anti-CD95 antibody Jo2 (BD Biosciences, 554254) or corresponding isotype control (BD Biosciences, 553961) was performed. Mice were culled 6 hours later and liver samples collected as described for experiment 1.

### Statistical analyses

All statistical analyses were performed using Graphpad PRISM (v9) and are described in the figure legends.

### RNAseq and TCGA analyses

For each sample, the XBP1s and RIDD activity scores were calculated using the 38 genes from the IRE1 signature^11^. The expression values of the 38 transcripts were extracted from the normalized count matrix of the 67 TNBC transcriptomes from a local cohort (GSE182021) and the 45 TCGA-GBM transcriptomes. Samples were then classified as high or low for XBP1s and RIDD activities according to the median. Then CD95, TNFR1, TRAIL-R1, TRAIL-R2, FADD or caspase-8 mRNA expression z-score was calculated for each sample and plotted according to the XBP1s and RIDD activities in both TNBC and GB datasets.

### Immunohistochemistry (IHC) and quantification

For IHC, samples were fixed in PFA 4% for at least 24 hours, embedded in paraffin at least 12 hours and sliced (4 μm) using a Leica microtome on Superfrost Plus slides (VWR, 631-0108) prior to drying at 60°C for 1 hour. The immunochemistry experiments were performed using the Discovery XT machine (Roche) and the Chromo-Map DAB kit (Roche). The following primary antibodies: cleaved caspase-3, Cell Signalling, 9661, diluted 1/300; CD95, R&D systems, AF435, diluted 1/50 in antibody diluent (NB-23-00171-1, NeoBiotech) were incubated for 1 hour at 37°C. To perform the analysis, glass slides were digitized with the scanner Nanozoomer 2.0-RS Hamamatsu. The quantifications of CD95 expression and cleaved caspase-3 staining were carried out using the NIS-Elements software (Nikon) and averaged from 5 fields covering 4 mm^2^ in total per liver.

## Acknowledgements

We thank all the members of the U1242 for fruitful scientific discussions. We thank Dr Marc Aubry for his help with bioinformatic analyses. We thank Drs Michel Samson and Jacques Le Seyec for their advice on the *in vivo* liver damage model. We thank the BIOSIT H2P2 platform for immunohistochemistry, in particular Gevorg Ghukasyan, and the BIOSIT Animal facility ARCHE (https://biosit.univ-rennes1.fr/). This work was funded by grants from Fondation ARC (PDF20171206671) and Fondation de France to EL, European Union (EU) H2020 MSCA ITN-675448 (TRAINERS) and INCa PLBIO 2018, 2019 grants to EC, PJA20181207700 (Fondation ARC) grant to EL and MLG, Ligue contre le cancer grants (from committees 22, 35, 36, 56, 85) to TA, Associations la Vannetaise et la Josselinaise des femmes funding to MLG and Centre Eugène Marquis (EL, EC, MLG, TA and SM).

## Author contributions

Conceptualisation: EC, EL; Formal analysis: AP, DPR, EL, MLG ; Funding acquisition: EC, EL, MLG; Investigation: AP, DPR, EL, RP, MLG, SM, TA, XZ ; Methodology: RP; Project administration: EL; Supervision: EC, EL; Validation: DPR, EL; Visualisation: AP, DPR, EL, MLG ; Writing-original draft: EL; Writing-review and editing: AP, TA, DPR, EC, EL, MLG, SM.

## References

1. Von Karstedt, S., Montinaro, A. & Walczak, H. Nature Reviews Cancer (2017).

2. Rossin, A., Miloro, G. & Hueber, A.O. Cancers 11 (2019).

3. Risso, V., Lafont, E., Le Gallo, M. Cell Death Disease 13, 248 (2022).

4. Almanza, A. et al. Febs j 286, 241–278 (2019).

5. McGrath, E.P., Centonze, F.G., Chevet, E., Avril, T. & Lafont, E. Biochimica Et Biophysica Acta Bba - Mol Cell Res, 119001 (2021).

6. Obacz, J. et al., 533018 (2020).

7. Doultsinos, D. et al. Biorxiv, 594630 (2019).

8. Drachsler, M. et al. Cell Death Dis 7, e2209 (2016).

9. Kleber, S. et al. Cancer Cell 13, 235–248 (2008).

10. Le Reste, P.J. et al., 841296 (2020).

11. Lhomond, S. et al. EMBO Mol Med 10 (2018).

12. Quijano-Rubio, C., Silginer, M. & Weller, M. Cell Death Discov 8, 341 (2022).

13. Chen, X. et al. Nature 508, 103–107 (2014).

14. Fritsche, H. et al. Oncotarget 6, 9502–9516 (2015).

15. Logue, S.E. et al. Nat Commun 9, 3267 (2018).

16. Harnoss, J.M. et al. Cancer Res 80, 2368–2379 (2020).

17. Maurel, M., Chevet, E., Tavernier, J. & Gerlo, S. Trends Biochem Sci 39, 245–254 (2014).

18. Lafont, E. Cancers 12, 1113 (2020).

19. Stöhr, D., Jeltsch, A. & Rehm, M. International review of cell and molecular biology 351, 57–99 (2020).

20. Li, T. et al. J Biol Chem 290, 11108–11118 (2015).

21. Lam, M., Lawrence, D.A., Ashkenazi, A. & Walter, P. Cell Death Differ 25, 1530–1531 (2018).

22. Lu, M. et al. Science 345, 98–101 (2014).

23. He, Q. et al. Oncogene 21, 2623–2633 (2002).

24. Yamaguchi, H. & Wang, H.G. J Biol Chem 279, 45495–45502 (2004).

25. Cazanave, S.C. et al. J Biol Chem 286, 39336–39348 (2011).

26. Jiang, C.C. et al. Cancer Res 67, 5880–5888 (2007).

27. Iurlaro, R. et al. Mol Cell Biol 37 (2017).

28. Iurlaro, R. & Munoz-Pinedo, C. Febs j 283, 2640–2652 (2016).

29. Lam, M., Marsters, S.A., Ashkenazi, A. & Walter, P. Elife 9 (2020).

30. Pluquet, O. et al. Cancer research 73, 4732–4743 (2013).

31. Nguyên, D.T. et al. Molecular biology of the cell 15, 4248–4260 (2004).

32. Avril, T. et al. Brain pathology (Zurich, Switzerland) 22, 159–174 (2012)

33. Volkmann, K. et al. J Biol Chem 286, 12743–12755 (2011).

34. Moore, K. & Hollien, J. Molecular biology of the cell 26, 2873–2884 (2015).

35. Oikawa, D., Tokuda, M., Hosoda, A. & Iwawaki, T. Nucleic Acids Res 38, 6265–6273 (2010).

36. Yoshida, H., Matsui, T., Yamamoto, A., Okada, T. & Mori, K. Cell 107, 881–891 (2001).

37. Voutetakis, K.D. D.; Vlachavas, E-I., Leonidas, DD.; Chevet, E.; Chatzioannou, A. (In preparation).

38. Geserick, P. et al. J Cell Biol 187, 1037–1054 (2009).

39. Jost, P.J. et al. Nature 460, 1035–1039 (2009).

40. Condon, S.M. et al. J Med Chem 57, 3666–3677 (2014).

41. Benetatos, C.A. et al. Molecular cancer therapeutics 13, 867–879 (2014).

42. Filliol, A. et al. Sci Rep 7, 9205 (2017).

43. Stöhr, D. et al. Cell Death Differ 27, 3037–3052 (2020).

44. Sullivan, G.P. et al. Dev Cell 52, 714-730.e715 (2020).

45. Hellwig, C.T. et al. Cell Death Differ 29, 147–155 (2022).

46. Timmins, J.M. et al. J Clin Invest 119, 2925–2941 (2009).

47. Miani, M., Colli, M.L., Ladrière, L., Cnop, M. & Eizirik, D.L. Endocrinology 153, 3017–3028 (2012).

48. Singh, N. et al. Cancer Discov 10, 552–567 (2020).

49. Upadhyay, R. et al. Cancer Discov 11, 599–613 (2021).

50. Martinez-Turtos, A. et al. Oncoimmunology 11, 2116844 (2022).

51. Annibaldi, A. & Walczak, H. Cold Spring Harb Perspect Biol 12, a036384 (2020).

52. Thamsen, M. et al. PLoS One 14, e0209824 (2019).

53. Auyeung, V.C. et al. American journal of physiology. Lung cellular and molecular physiology (2022).

54. Redente, E.F. et al. JCI insight 6 (2020).

55. Guégan, J.P. et al. iScience 24, 103538 (2021).

56. Risso, V. et al. FEBS J (In press) (2022).

57. Lorenz, R. et al. Algorithms for molecular biology : AMB 6, 26 (2011).

58. Schneider, C.A., Rasband, W.S. & Eliceiri, K.W. Nat Methods 9, 671–675 (2012).

