## Supplementary data for "IRE1 RNase controls CD95-mediated cell death"

**IRE1 RNase controls CD95-mediated cell death- Supplementary Figures**

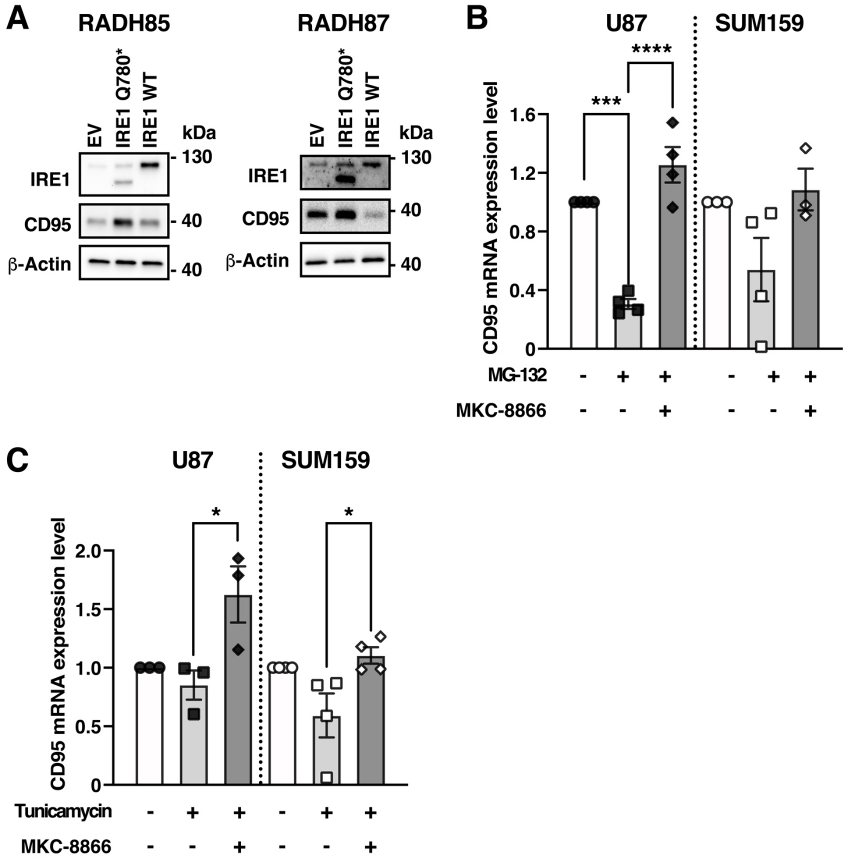


**Supplementary Figure 1 - IRE1 represses CD95 expression in GB cells upon ER stress. A.** CD95 protein level was evaluated by western blot in lysates from the indicated cells. One representative experiment out of three independent experiments is presented. **B.** U87 or SUM159 cells were pre-incubated for 1 hour with 1μg/mL of actinomycin D and further treated with 10μM MKC-8866 for 1 hour followed by 2 hours treatment with 10μM MG-132 as indicated. **C.** U87 or SUM159 cells were pre-incubated for 1 hour with 1μg/mL of actinomycin D and further treated with 10μM MKC-8866 for 1 hour followed by 2 hours treatment with 1μg/mL tunicamycin as indicated. **B, C**: Mean ± SEM, n=3-4. *p<0.05, ***p<0.001, **** p<0.0001, one-way ANOVA with Dunnett multiple comparison correction.

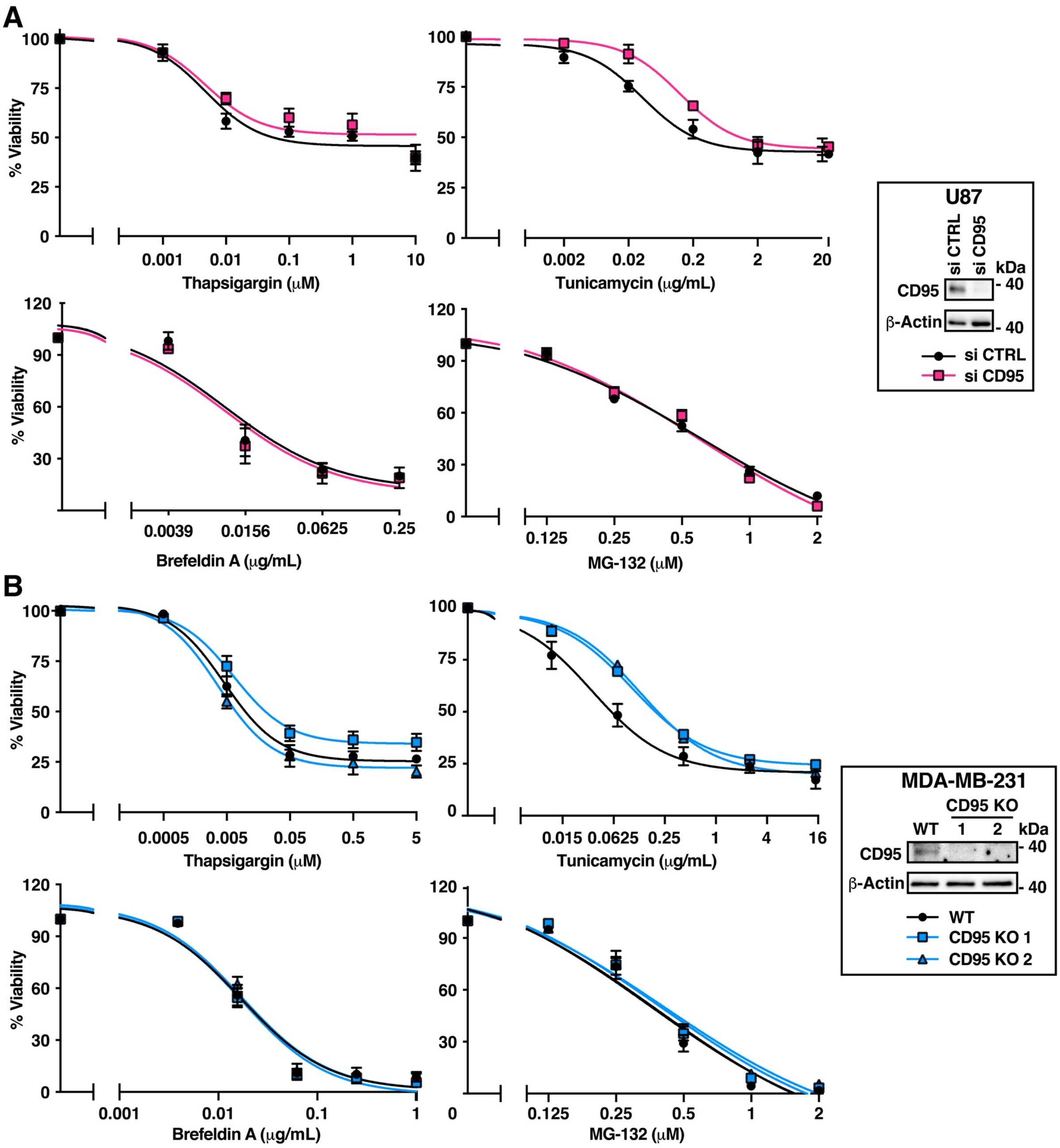


**Supplementary Figure 2 - CD95 is not a general determinant of ER stress-induced cell death. A.** U87 transfected with control (CTRL) or CD95-targeting siRNAs were treated with the indicated ER stress inducers for 48 hours. Viability was assessed using MTT assay. Mean ± SEM, n=3-4 independent experiments. **B.** MDA-MB-231 WT or CD95 KO clones were treated with the indicated ER stress inducers for 48 hours. Viability was assessed using MTT assay. Mean ± SEM, n=4-5 independent experiments. **A, B.** Insets: lysates were analysed by western blot using the indicated antibodies.

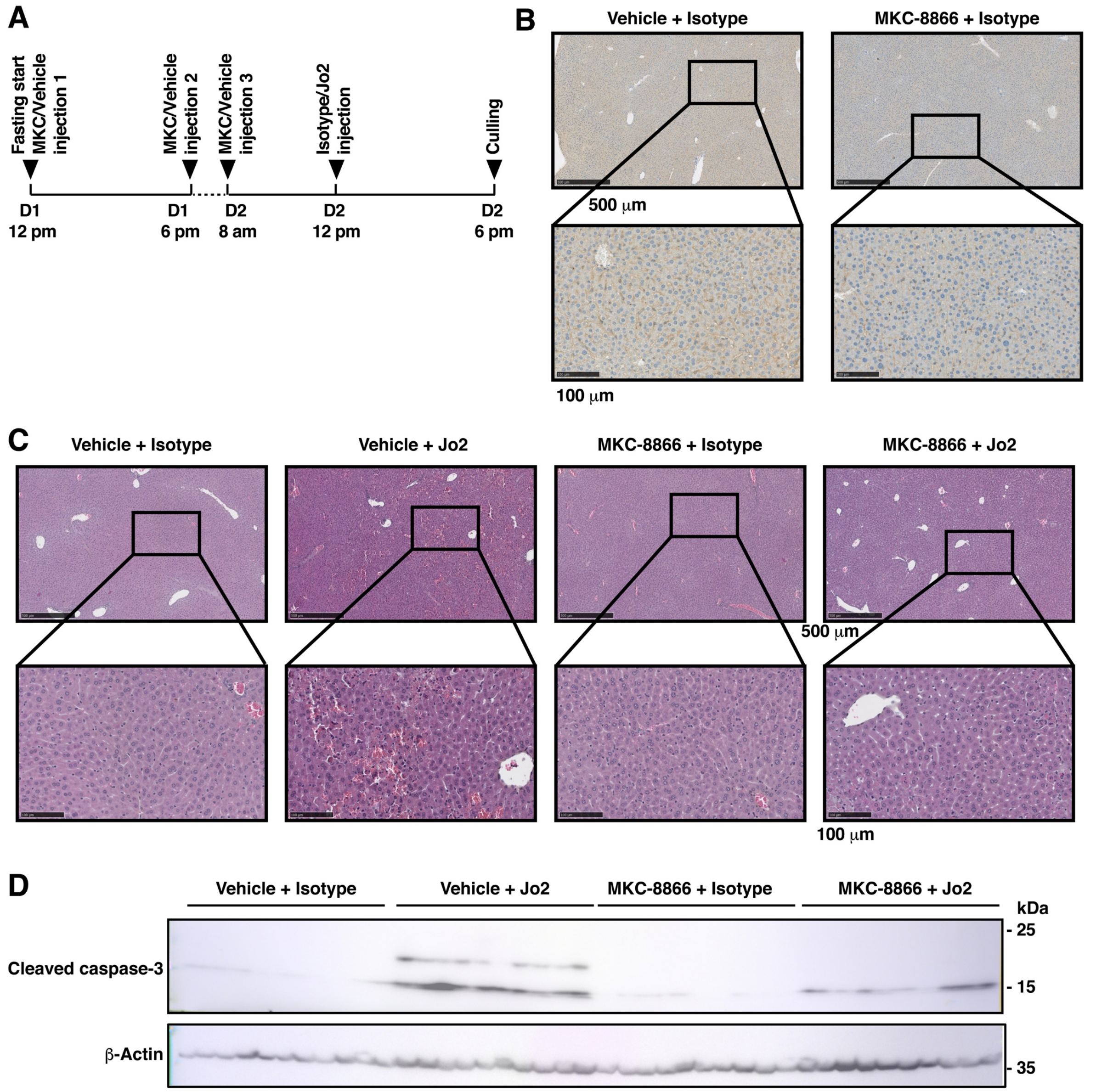


**Supplementary Figure 3 - IRE1 RNase inhibition limits hepatic CD95 expression and CD95-mediated cell death in mice. A.** Timeline of the second *in vivo* experiment. 40 mice were divided in two groups of 20 and were repeatedly injected with either vehicle or MKC-8866 as indicated. On day 2 at 12pm, each of this initial groups were further divided in two groups of 10 mice which were injected with either an anti-CD95 antibody or with an isotype control as indicated. **B.** CD95 expression was evaluated by IHC in mice injected with vehicle or MKC-8866 and the isotype control antibody. One representative image is shown for each of these two groups. **C.** HES staining was performed on liver tissue sections from mice of each of the 4 groups described in A. One representative image is shown for each of these groups. **D.** Western blot analysis of liver lysates from mice treated as indicated.

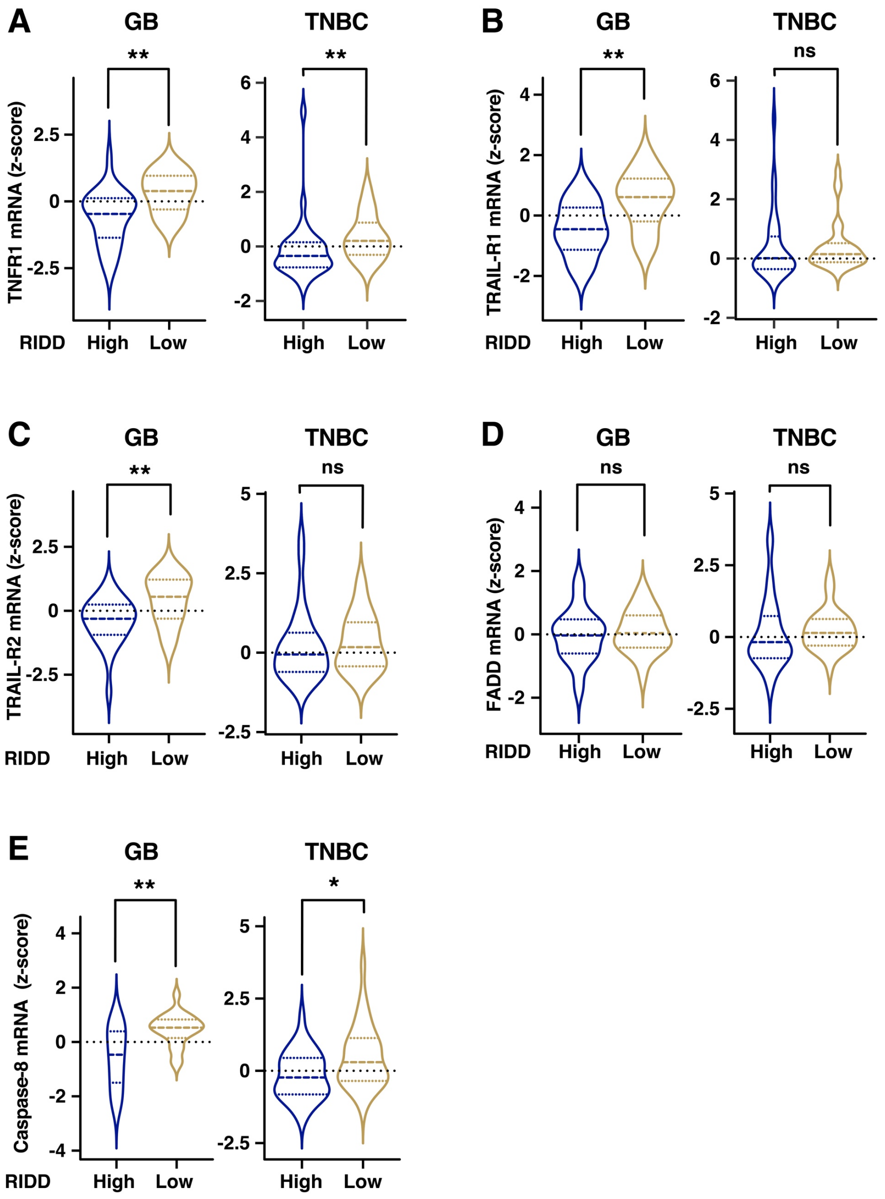


**Supplementary Figure 4 - Low RIDD activity in tumors correlates with the expression of core pro-apoptotic DR signaling components.** TNFR1 (**A**), TRAIL-R1 (**B**), TRAIL-R2 (**C**), FADD (**D**) or caspase-8 (**E**) expression z-scores of 45 GB and 67 TNBC tumors were plotted according to the RIDD activity score. The distribution of z-score is represented as violin plots. Statistical difference of expression between groups was calculated using Mann-Whitney tests and the p-value is indicated (*<0.05, ** p<0.01, ns: not significant).

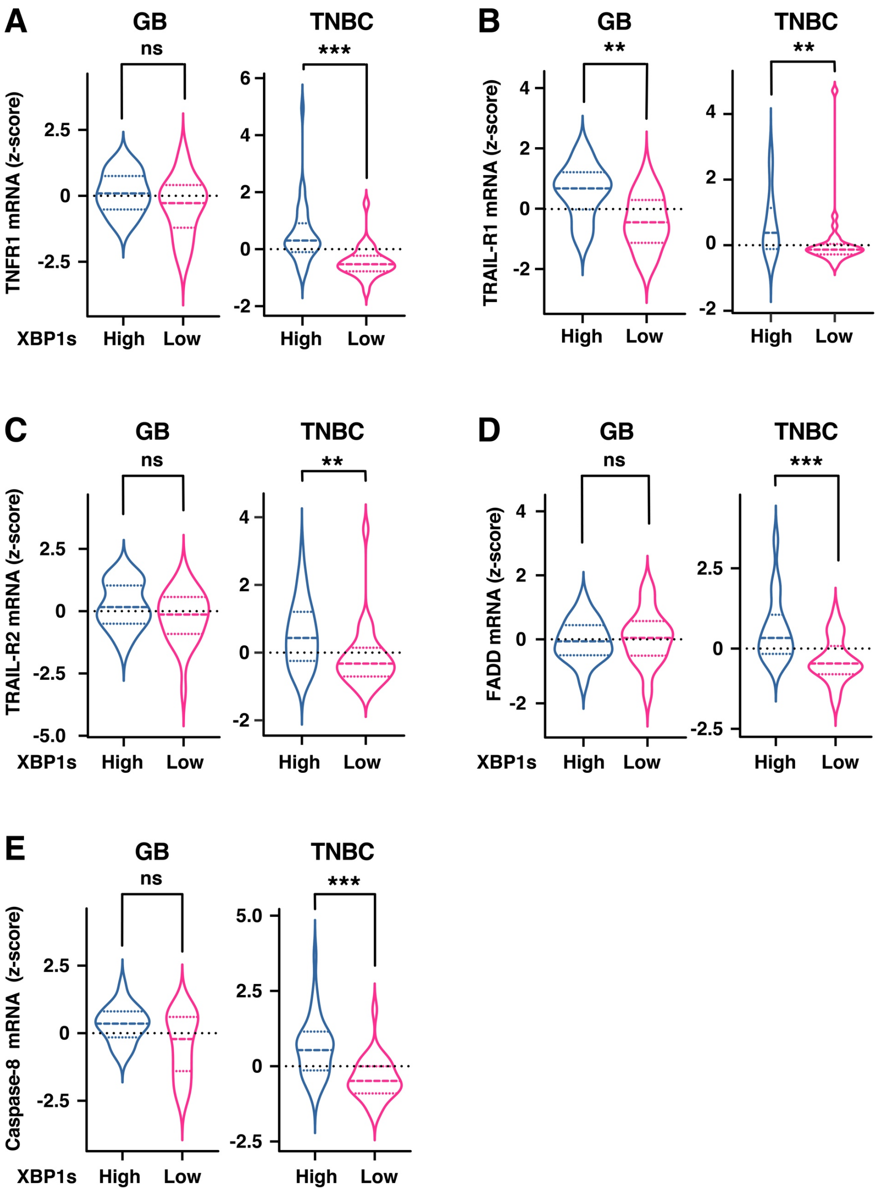


**Supplementary Figure 5 - High XBP1s activity in tumors correlates with the expression of core pro-apoptotic DR signaling components.** TNFR1 (**A**), TRAIL-R1 (**B**), TRAIL-R2 (**C**), FADD (**D**) or caspase-8 (**E**) expression z-scores of 45 GB and 67 TNBC tumors were plotted according to the XBP1s activity score. The distribution of z-score is represented as violin plots. Statistical difference of expression between groups was calculated using Mann-Whitney tests and the p-value is indicated (** $p<0.01$, *** $p \leq 0.001$, (*<0.05, ns: not significant)).
